# *Cryptococcus neoformans* releases proteins during intracellular residence that affect the outcome of the fungal-macrophage interaction

**DOI:** 10.1101/2021.09.21.461206

**Authors:** Eric H. Jung, Yoon-Dong Park, Quigly Dragotakes, Lia Sanchez Ramirez, Daniel Q. Smith, Amanda Dziedzic, Peter R. Williamson, Anne Jedlicka, Arturo Casadevall, Carolina Coelho

**Affiliations:** Department of Molecular Microbiology and Immunology, Johns Hopkins School of Public Health, Baltimore, Maryland, USA; Laboratory of Clinical Immunology and Microbiology, National Institute of Allergy and Infectious Disease, National Institutes of Health, Bethesda, Maryland, USA; Department of Molecular and Cell Biology, Johns Hopkins University, Baltimore, Maryland, USA; MRC Centre for Medical Mycology, College of Health and Medicine, University of Exeter, Exeter, Devon, UK

## Abstract

*Cryptococcus neoformans* is a facultative intracellular pathogen that can replicate and disseminate in mammalian macrophages. In this study, we analyzed fungal proteins identified in murine macrophage-like cells after infection with *C. neoformans*. To accomplish this, we developed a protocol to identify proteins released from cryptococcal cells inside macrophage-like cells; we identified 127 proteins of fungal origin in infected macrophage-like cells. Among the proteins identified was urease, a known virulence factor, and others such as transaldolase and phospholipase D, which have catalytic activities that could contribute to virulence. This method provides a straightforward methodology to study host-pathogen interactions. We chose to study further Yor1, a relatively uncharacterized protein belonging to the large family of ATP binding cassette transporter (ABC transporters). These transporters belong to a large and ancient protein family found in all extant phyla. While ABC transporters have an enormous diversity of functions across varied species, in pathogenic fungi they are better studied as drug efflux pumps. Analysis of *C. neoformans yor1*Δ strains revealed defects in non-lytic exocytosis and capsule size, when compared to wild-type strains. We detected no difference in growth rates, cell body size and vesicle secretion. Our results indicate that *C. neoformans* releases a large suite of proteins during macrophage infection, some of which can modulate fungal virulence and are likely to affect the fungal-macrophage interaction.

## Introduction

*Cryptococcus neoformans* is a basidiomycetous opportunistic fungal pathogen found worldwide (Casadevall and Perfect, 1998). Human infection occurs when desiccated yeasts or spores are inhaled into the lung where the infection is controlled in the majority of immunologically intact individuals (May et al., 2016). However, in many immunocompromised individuals, such as patients with HIV/AIDS or those on an immunosuppressive therapy, the infection is no longer controlled and cryptococcosis ensues, which often manifests itself as a subacute meningoencephalitis that is inevitably fatal if not treated with antifungal agents. While this disease predominantly affects immunocompromised individuals, it occurs sporadically in immunocompetent individuals (Fisher et al., 2016). Cryptococcosis results in approximately 180,000 deaths per annum, primarily in sub Saharan Africa where HIV/AIDS prevalence is high (Rajasingham et al., 2017).

*C. neoformans* is a facultative intracellular pathogen that can replicate inside macrophages (Feldmesser et al., 2000) and the topic has been the subject of several recent reviews (Coelho et al., 2014; DeLeon-Rodriguez and Casadevall, 2016; Gilbert et al., 2014; Mansour et al., 2014). There is a correlation between the susceptibility of mice and rats to cryptococcosis and the ability of *C. neoformans* to replicate within macrophages (Shao et al., 2005; Zaragoza et al., 2007). In both mice and zebrafish increases in *C. neoformans* numbers are associated with the ability of fungal cells to replicate in macrophages (Bojarczuk et al., 2016; Feldmesser et al., 2000). In humans, there is a correlation between disease outcome and the intracellular replication of *C. neoformans* (Alanio et al., 2011). *C. neoformans* is able to survive and even thrive in the acidic and proteolytic phagolysosome, its primary location when ingested by host macrophages (De Leon Rodriguez et al., 2018; Fu et al., 2018). The three potential outcomes of the ingestion of *C. neoformans* by host macrophages are host killing of the fungal pathogen, intracellular fungal replication or egress of the fungi from the host macrophage by lytic or non-lytic exocytosis (DeLeon-Rodriguez and Casadevall, 2016; Zhang et al., 2015). Non-lytic exocytosis is the ability of the yeast to escape from its host macrophage with both host and pathogen remaining viable (Alvarez and Casadevall, 2006; Ma et al., 2006) and this phenomenon has been shown to occur *in vivo* in mice (Nicola et al., 2011) and zebra fish (Bojarczuk et al., 2016). Initially, the three types of non-lytic exocytosis were classified as: type I, the complete extrusion of fungal burden from the host macrophage; type II, the partial expulsion of the fungal burden with at least one yeast remaining; and type III, the cell-to-cell transfer of one or more *C. neoformans* cells between host macrophages (Stukes et al., 2016, 2014). However, recent work has established that cell-to-cell transfer is the result of a sequential non-lytic exocytosis event followed by phagocytosis of an adjoining cell, a process termed ‘Dragotcytosis’ (Dragotakes et al., 2019). Thus, the mechanisms for non-lytic exocytosis remain poorly understood, however there has been an increasing body of evidence that it may be an important factor in *C. neoformans*-macrophage interaction.

There is considerable evidence that intracellular *C. neoformans* residency actively modulates the physiology of macrophages. In this regard, early studies showed that polysaccharide-laden vesicles that appeared to bud from cryptococcal phagosomes accumulated in the macrophage cytoplasm (Feldmesser et al., 2000). Phagosomal membranes become leaky with time (De Leon Rodriguez et al., 2018; Tucker and Casadevall, 2002), reflecting damage from secretion of phospholipases and enlargement of the capsule (De Leon Rodriguez et al., 2018). Intracellular residence is associated with impairment of mitochondrial function and activation of cell death pathways (Coelho et al., 2015). *C. neoformans* affects host cell NF-κB expression (Hayes et al., 2016), actin cytoskeleton dynamics (Johnston and May, 2010), phagosomal pH (Dragotakes et al., 2020; Fu et al., 2018) and gene transcription profiles(Subramani et al., 2020) through mechanisms that are poorly understood, but are likely to reflect effects of cryptococcal products which interfere with host cell physiology. In this regard, *C. neoformans* has recently been shown to release small molecules that affect macrophage function (Bürgel et al., 2020). *C. neoformans* is known to release numerous proteins into the exterior of the cell (Chen et al., 1996; Geddes et al., 2015), some of which are released in extracellular vesicles (Rodrigues et al., 2008a). The mechanisms by which *C. neoformans* modulates and damages macrophages are poorly understood, but are likely to reflect effects of cryptococcal products on host cell physiology. In this study, we investigated whether *C. neoformans* released proteins into macrophage-like J774.1 immortalized cells. The results show that *C. neoformans* infection of macrophage-like cells is followed by the production of numerous proteins, including a <u>Y</u>east <u>O</u>ligomycin <u>R</u>esistance (Yor1) protein, an ABC transporter whose deletion affects the frequency of non-lytic exocytosis.

## Results

### Approaches to study *C. neoformans* proteins secreted during infection of murine macrophages

To demonstrate the presence of cryptococcal proteins in macrophages, the first approach we attempted was to radioactively label the proteins of *C. neoformans*, then add these labelled-fungi to macrophages for phagocytosis and analyze whole co-infection lysates for the presence of radioactive proteins. Hence, we grew *C. neoformans* with different concentrations of ^35^S-methionine and ^35^S-cysteine. We observed normal growth curves in different concentrations of radiolabeling material (Figure 1A). Next, we analyzed the labeling efficiency of *C. neoformans* and determined that H99 strain grown in the presence of 20 µCi ^35^S-Met and ^35^S-Cys during a 4-hour incubation yielded the greatest labeling efficiency (Figure 1B). We then infected J774.16 macrophages with radiolabeled *C. neoformans*, lysed the infected murine cells and subsequently run whole cell lysates on a 2-D isoelectric focusing apparatus. This revealed that detectable amounts of radiolabeled-fungal proteins were released into the J774.16 macrophages. Further, the protein profile was different from that observed in a lysate of *C. neoformans* grown in Sabouraud media (Figure 1C). Having established that fungal proteins can be identified from co-cultures of *C. neoformans* infected J774.16-macrophages, and with the goal of identifying the secreted fungal proteins via proteomic mass spectroscopy, we attempted nonradioactive methods of protein labeling. First, we attempted to label newly-synthesized cryptococcal proteins by growing fungi in the presence of tRNA pre-charged with biotinylated lysine, but observed no cryptococcal labeling (data not shown). Second, we attempted click-chemistry by labeling nascent proteins with L-Homopropargylglycine (HPG), a glycine analogue containing alkyne moiety that can be subsequently click-labelled. Again, we were unable to observe detectable labelled-protein bands, despite the fact that the mammalian Jurkat cells, used as a positive control for click-chemistry, did yield distinct, click-labeled protein (data not shown). We attempted two other methods to characterize the cryptococcal proteins released intro macrophages, namely to label proteins using a tRNA charged with biotinylated lysine, and fungal-phagosome isolation, but none was successful. Although we do not have explanation as to why tRNA charging and click chemistry failed, we suspect that these reflect peculiarities specific to the cryptococcal system. The failure to isolate intact *C. neoformans*-containing phagosomes by the well-tried magnetic bead technique, so successful when applied to bacterial phagosomes, may simply reflect the large size of the cryptococcal phagosome, and the fact that *C. neoformans* phagosomal membranes are damaged during the intracellular infection process (Tucker and Casadevall, 2002). The large size of the phagosome can leave the membrane vulnerable to shear forces and accentuated by *C. neoformans*-driven membrane damage (De Leon Rodriguez et al., 2018). We decided to resort to a selective extraction protocol.

**Figure 1.**
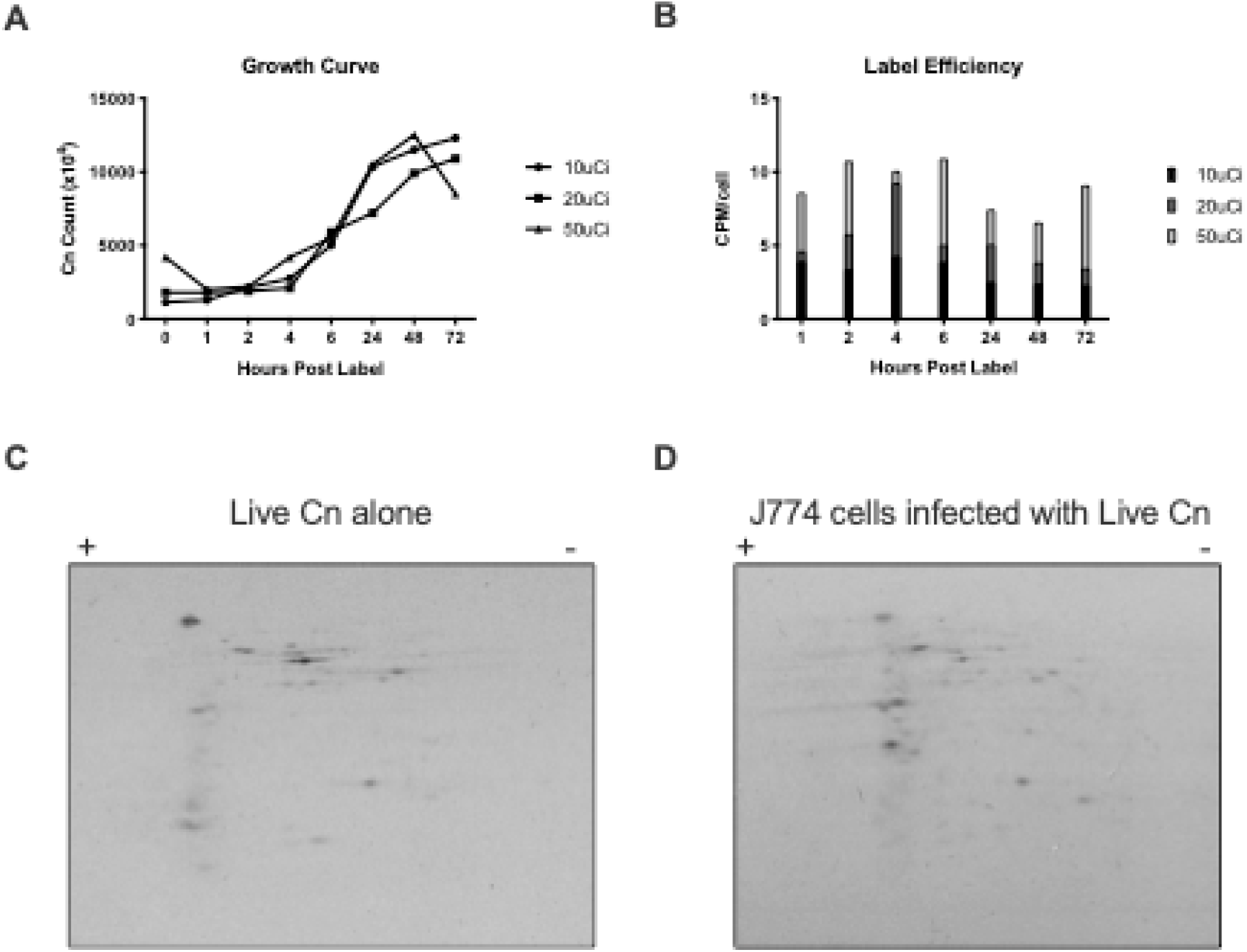
Radiolabeling *C. neoformans* proteins to detect putatively secreted proteins during mammalian infection. (A)^35^S labeling does not affect growth of *C. neoformans*. Growth was measured after addition of ^35^S-methionine and -cysteine to Sabouraud broth, at 30° C with constant agitation, at different time points over 72 h. (B) *C. neoformans* incorporates ^35^S-aminoacids. CPM measurements were made for each of the ^35^S labeling conditions by sampling at each time point. to calculate CPM/cell was calculated by (total CPM)/(total cell count). Shown is the mean CPM/cell for each time point. We determined 20 μCi and 4 h of incubation to yield maximum label and minimizing excess radiation for subsequent experiments. (C and D) As a proof of concept for efficient cryptococcal labeling and identification of putatively secreted proteins during infection, we labelled *C. neoformans* with ^35^S, and prepared protein lysates from broth culture and from infected J774.16 murine macrophage-like cells, followed by extraction of proteins from the co-culture and then run in a 2-D gel. Experiment shows the opportunity for isotopic labeling of aminoacids to detect differentially expressed proteins during host-pathogen interactions.

### *C. neoformans* protein profile

Given that the capsule and cell wall confer upon fungal cells a tremendous structural strength that requires harsh methods for extracting proteins relative to those required for lysing macrophage-like cells, we reasoned that we could lyse and extract proteins from the host cells while leaving internalized *C. neoformans* cells intact. Consequently, we devised and tested a protein extraction protocol designed to extract all host J774.16 proteins which would not lyse and extract fungal intracellular proteins. This protocol would allow proteins secreted by *C. neoformans* to be co-extracted within the mammalian cell lysates and their identity distinguishable by computational proteomic analysis. First, we subjected live *C. neoformans* cells in the absence of macrophages to a widely used mammalian lysis process which consists of multiple rounds of passaging through a 26 G needle; this process yielded no identifiable cryptococcal proteins. Hence, the isolation process used did not release proteins from *C. neoformans* viable cells. Having established a lysis and proteomics protocol, we infected J774.16 macrophages with live fungi from an H99 fungal strain (live H99); as a control, J774.16 macrophages were infected with heat-killed H99 (HK H99), reasoning that no active secretion is occurring in HK-H99 and the identified proteins would instead derive from macrophage-degradation of fungal remnants. We identified 127 cryptococcal proteins in lysates of J774.16-macrophages infected live H99 and 117 proteins from macrophages infected with HK H99. The cryptococcal proteins extracted from macrophage-like cells containing live compared to heat-killed *C. neoformans* had only 8% (18/226 proteins) proteins in common (Figure 2, Table 1, Supplemental Table 1), supporting our rationale. Among the 109 proteins identified in lysates of macrophages that had ingested live *C. neoformans* was urease, a known secreted virulence factor, among others such as transaldolase and phospholipase D, which are less characterized but could conceivably be secreted. To extract information on these proteins, we took advantage of available bioinformatic analysis to predict transmembrane domains and signal sequences. Of these fungal proteins, 19 proteins are predicted to have a signal sequence according to at least one of the three tools and 39 have at least one transmembrane domain (Supplemental Table 1). For the purposes of this report we did not analyze the host proteins.

**Table 1.** Top-ten *C. neoformans* proteins identified in J774.16 macrophage infected with live H99.

| Gene | Protein (length) | Molecular Weight (kDa) | Spectral count | Features | Comments |
| --- | --- | --- | --- | --- | --- |
| CNAG_01984 | transaldolase (324 aa) | 35 kDa | 24.407 |  | Role in CN virulence is unknown. Involved in resistance to nitric oxide (NO)(Missall et al., 2006), which may aid CN intracellular survival. |
| CNAG_06812 | phospholipase D (1745 aa) | 196 kDa | 13.054 |  | Role in CN virulence is unknown. Unlike Plp1, not detected as secreted in culture (Chen et al., 1997) |
| CNAG_05894 | dynein heavy chain 1, cytosolic (4630 aa) | 524 kDa | 10.331 |  | Transport (via GO) |
| CNAG_03655 | hypothetical protein (2007 aa) | 222 kDa | 9.2767 |  | 43% homology to *S. cerevisiae* *Num1p*, which is involved in organelle migration |
| CNAG_04172 | transcription factor C subunit 6 (826 aa) | 91 kDa | 6.7986 |  | Transcription/Translation |
| CNAG_02103 | hypothetical protein (1896 aa) | 209 kDa | 6.4459 |  | Homologs in other fungi but no function prediction |
| CNAG_00583 | hypothetical protein (1109 aa) | 118 kDa | 6.4459 | 1 TM | No significant homology detected |
| CNAG_06332 | hypothetical protein (1093 aa) | 121 kDa | 5.8274 |  | No significant homology detected, increased virulence(Gerstein et al., 2019) |
| CNAG_06728 | kinesin (958 aa) | 107 kDa | 4.8814 |  | Transport |
| CNAG_06818 | hypothetical protein (837 aa) | 93 kDa | 4.2973 |  | likely homology to Hap1 of *Saccharomyces cerevisiae* (Jung et al., 2010) |
| CNAG_07781 | ATP-dependent bile acid transporter (1618 aa) | 180 kDa | 4.2973 | 13 TM | Transport |
| CNAG_04370 | U3 small nucleolar RNA-associated protein 10 (2022 aa) | 222 kDa | 4.2973 | SP | Ribosomal Processing |

TM=transmembrane domain predicted, SP=signal peptide predicted

**Figure 2.**
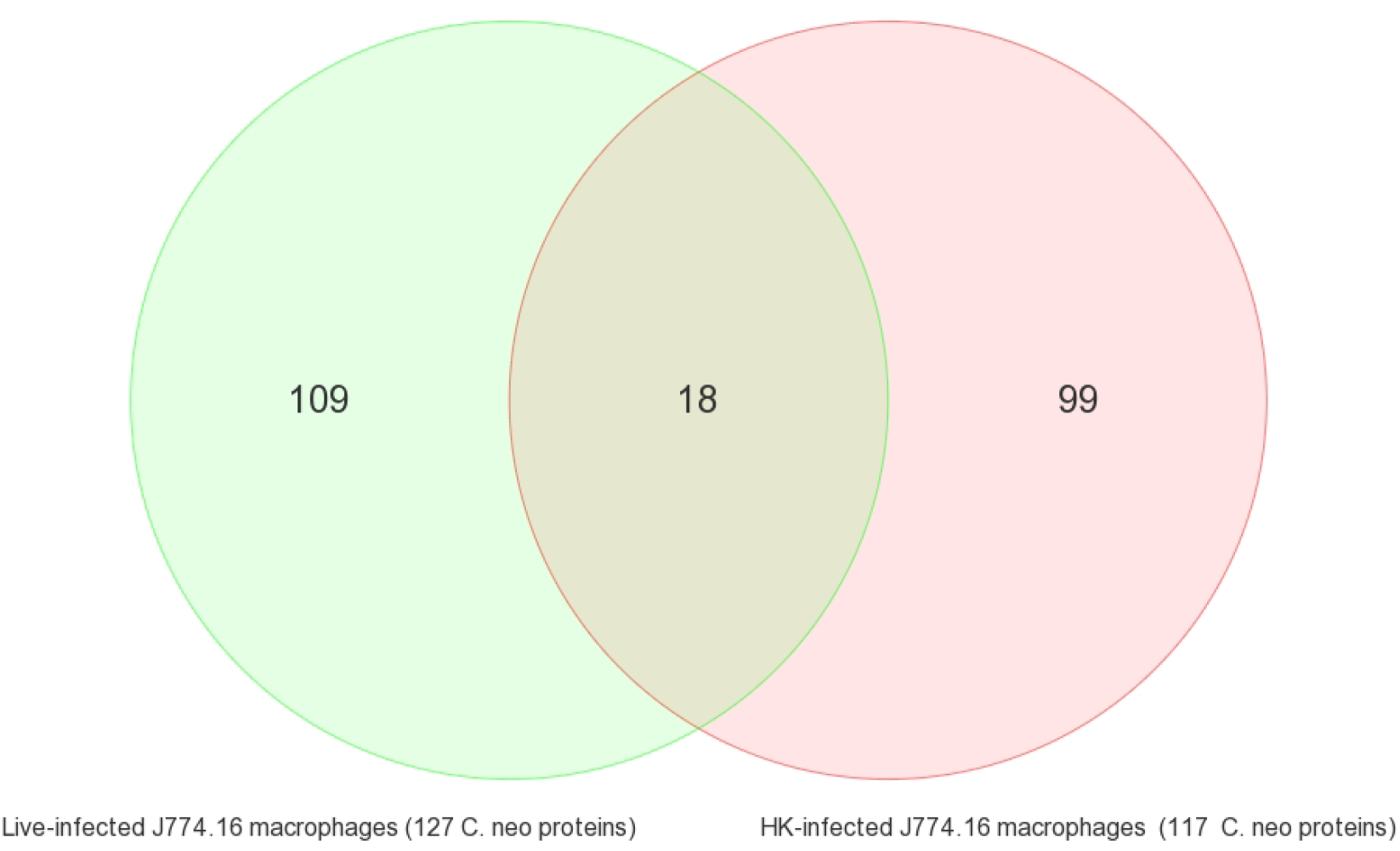
Proteomics analysis detected different cryptococcal proteins when J774.16 cells were infected with live Cn versus HK Cn. Using spectral count as a method of semi-quantitative expression levels, we identified 127 from live *C. neoformans* and 117 cryptococcal proteins from heat-killed (HK) *C. neoformans*, with little overlap between identified proteins. Full list of proteins in Supplemental Table 1.

### Identification of new virulence factors in *C. neoformans*

To validate that our strategy could identify novel fungal virulence factors, we then performed a preliminary analysis of putative novel virulence factors. Among the proteins identified macrophage-like cells infected with live *C. neoformans* was the ABC transporter Yor1, with thus far unknown roles in virulence. To study its role in fungal biology and virulence, and validate our screening strategy, we studied both a deletion strain from an available deletion library, generated by Madhani laboratory (*yor1*Δ), as well as generated *de novo Yor1*-deleted strains of *C. neoformans*, resorting to well-established replacement with *URA5*-selection cassette and biolistic particle delivery to remove the coding sequence of *Yor1* (Supplementary Figure 1). Screening with colony-PCR identified three independent *yor1*Δ strains (*yor1*Δa-c). For the *de novo* deletion strains generated, we characterized virulence factors, such as melanin production, virulence in *Galleria melonella*, capsule size in minimal media (MM), cell body size, capacity to alter phagosomal pH, and growth rates (Supplemental Figure 2-5). To evaluate whether *YOR1* deletion was associated with decreased melanization, we grew the *C. neoformans yor1Δ* strain from the 2008 Madhani knockout library, the three independently generated *yor1Δa, yor1Δb*, and *yor1Δc*, and their parental strains H99W and H99, respectively, in minimal media containing L-DOPA. These growth conditions normally induce melanisation (Supplemental Figure 2). Following three days of growth at 30°C, there is a notable defect in melanization in both sets of *yor1Δ* mutants, with the notable exception of *yor1Δc* which exhibits wild type melanisation.

To assay whether the *Yor1* was involved in virulence of *C. neoformans*, we used the *Galleria mellonella* wax moth model of infection (Supplementary Figure 3). We observed that *yor1Δ* strain from the 2008 strain library had reduced virulence compared to the parental strain from the library (H99W). For *yor1*Δa and *yor1*Δb, there was a similar reduction in virulence. Strain *yor1*Δc is displaying different phenotypes than its counterparts *yor1*Δa and *yor1*Δb, which may be due to spurious secondary mutations in this strain, albeit we we did not investigate this further. Overall, in 3 of 4 deletion strains Yor1-deletion was associated with reduced melanization and virulence in wax moth.

We detected a reduction in capsule size in *yor1*Δ strains in minimal media, but this was dependent on parental background (Supplemental Figure 4). Thus, it is possible that Yor1 is involved in capsular synthesis, secretion or assembly, albeit this is dependent on strain background. We had previously associated capsule changes and urease production with capacity to manipulate phagolysosomal pH(De Leon-Rodriguez et al., 2018, p.; Fu et al., 2018). We found that Yor1 is not involved in manipulating phagolysosomal pH. To assess any growth defects in *yor1*Δ, we performed growth curves in liquid media over the course of 72-96 h(Supplementary Figure 5). When analyzing cultures grown in Sabouraud or MM, we observed no growth defects: a similar lag phase duration, slope of log phase, and establishment of stationary phase.

We also evaluated rates of non-lytic exocytosis *in vitro* of *yor1*Δ strains when exposed to bone marrow-derived macrophages (BMDM)s. J774.16 cells were not used as their high motility is challenging for non-lytic exocytosis quantification. We infected BMDMs with *yor1*Δ strains and wild-type H99 (Figure 3) and measured non-lytic exocytosis rates in three, independent 24 h time-lapse movies with BMDMs infected with wild type H99 with 50 to 100 cells tracked per individual movie. We observed a mean non-lytic exocytosis rate reduction in all *yor1*Δ strains. The different parental backgrounds of deletion strains may generate some variability, but overall the defects are the same. Although we were not able to complement the *yor1*Δ strains, we note that the reduced non-lytic exocytosis phenotype was observed with four independent mutants, which provides confidence for a causal association. A search in available data, via searches in FungiDB and ClustalOmega, showed that other strains of C. neoformans have Yor1, with a close homologue (CNBG_2112) in *C. gattii* strain R265. These searches also identified that *Yor1* is upregulated after incubation in DMEM, 5% CO_2_. To assess the integrity of the library strains for a non-specific defect that affected non-lytic exocytosis, and thus if genetic manipulation of fungi could affect non-lytic exocytosis, we analyzed the *cir1*Δ (CNAG_04864), a strain known to have defects in iron acquisition which causes defects in virulence (Jung et al., 2006), and ascertained that despite known defects in virulence, the rate of non-lytic exocytosis of *cir1*Δ is comparable to that of the wild type H99 strain (data not shown).

**Figure 3.**
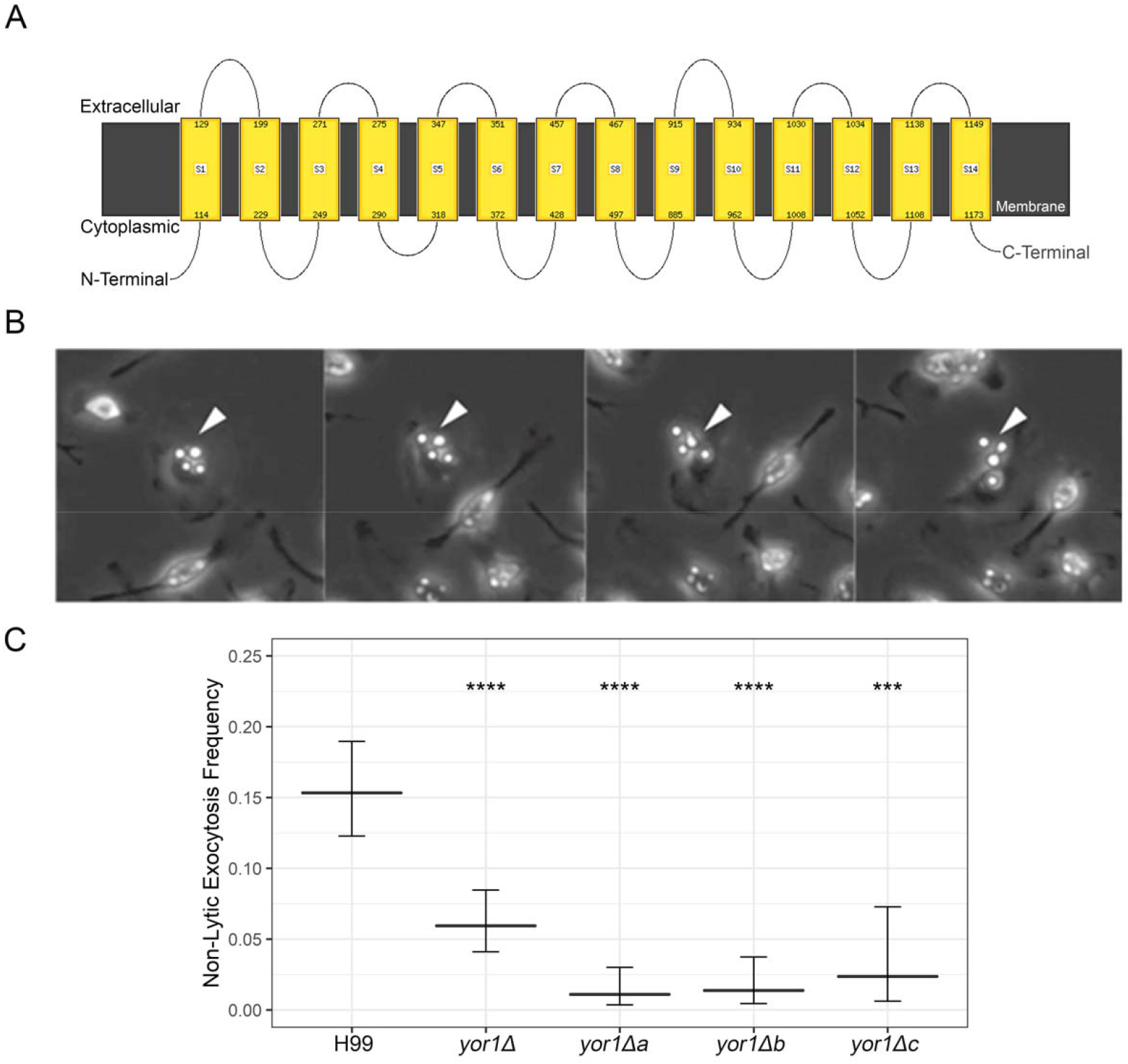
Yor1, a transmembrane protein, affects non-lytic exocytosis of *C. neoformans*. A) Schematic of Yor1, showing transmembrane domains, as predicted by bioinformatic tool Phyre2. **(A)** H99 and *yor1*Δ-strains were opsonized with mAb 18B7 prior to phagocytosis by 7 d old BMDMs from C57Bl/6 mice. After time lapse microscopy at 10× magnification for 24 h, each macrophage was tracked for non-lytic exocytosis (white arrows). **(B)** Strains with *yor1*Δ as well as each of three de novo mutant strains show a defect in non-lytic exocytosis compared to wildtype strain H99. Comparisons were performed with a two tailed test of equal proportions compared to the wild type H99 strain with Bonferroni correction for multiple tests. *** and **** signify *p* < 0.001 and 0.0001, respectively. Error bars denote 95% confidence intervals.

## Discussion

This study investigated *C. neoformans* proteins in macrophages after fungal cell ingestion in macrophage-like cells. The study was prompted by recent evidence that intracellular residence of *C. neoformans* in macrophage-like cells is associated with changes to host cell physiology and metabolism (Coelho et al., 2015), which presumably reflect host cell damage and/or modulation by the fungal cells. Proteins secreted by human pathogenic fungi during macrophage interaction *in vivo* has been determined before for *Candida albicans* (Kitahara et al., 2015) and *Aspergillus fumigatus* (Schmidt et al., 2018), but not for the *C. neoformans* species complex. We focused on developing a protocol to identify putatively secreted proteins, as those may have direct effects in modulating host cells and are usually major virulence factors and/or immunogens. The major finding in our study is that *C. neoformans* intracellular residence is associated with the production of a variety of protein products and some of these proteins have important roles in virulence and pathogenesis.

*C. neoformans* protein secretion in macrophage-like cells was established by two independent approaches. First, we show, by two-dimensional protein electrophoresis after phagocytosis of ^35^S-labelled *C. neoformans* cells, that a new set of fungal proteins is discernible in host cells lysates, presumably by secretion of fungal proteins during mammalian infection (Figure 1C-D). Second, we used mass spectroscopy to identify fungal proteins co-isolated in macrophage lysates that ingested live or killed *C. neoformans* to identify the secreted proteins of *C. neoformans*.

The protein sets identified in J774.16 macrophage-like cells that ingested live or dead *C. neoformans* cells were different. This provides strong support for the hypothesis that the proteins isolated from macrophage-like cells that ingested live *C. neoformans* cells were produced and secreted by live fungal cells while inside the macrophage and was not the result of macrophage digestion of fungi. Regardless of secretion, the co-isolation and identification in mammalian lysates, strongly indicates these proteins are produced in high levels *in vitro* during mammalian infection. These fungal proteins identified within infected macrophage-like cells could be located at the cryptococcal phagosomes and/or the cytoplasm, given that cryptococcal-phagosomes macrophages are leaky (De Leon Rodriguez et al., 2018; Tucker and Casadevall, 2002). We found no overlap with proteins previously identified in cryptococcal secreted proteins (Geddes et al., 2015) or associated with extracellular vesicles (Rodrigues et al., 2008b). Given that these studies were performed by growing fungi in microbiological media and that our study was performed on fungal-infected murine macrophages, we reason that these differences are likely due to these very distinct environmental conditions. Cryptococcal proteins isolated from macrophage-like cells containing live *C. neoformans* cells included some the well-known virulence factors such as urease Ure1 (Cox et al., 2000; Fu et al., 2018; Singh et al., 2013), as well as Zsd3 (Li et al., 2011), but the large majority of proteins identified were either previously unimplicated in virulence or hypothetical proteins due to significant divergence to known proteins. Urease is required for brain dissemination of *C. neoformans* and well-known for being secreted (Cox et al., 2000; Fu et al., 2018; Singh et al., 2013). Urease activity prevents acidification of fungal-containing phagolysosomes, and is required for optimal growth at mammalian pH (Fu et al., 2018). We also identified phospholipase D (PLD). Whereas phospholipase B is a well-characterized virulence factor in *C. neoformans* pathogenicity (Cox et al., 2001; Djordjevic, 2010; Noverr et al., 2003, p. 1) that was recently implicated in mediating damage to the membrane of phagolysosomes containing *C. neoformans* (De Leon Rodriguez et al., 2018), PLD has not been implicated in virulence thus far. Previous analysis of *C. neoformans* supernatants obtained in microbiological media supplemented with egg yolk revealed no PLD activity (Chen et al., 1997). It is interesting that the gene encoding for PLD, CNAG_06812, is located in the alpha mating locus, which is associated with virulence (Sun et al., 2020). Expression of this protein inside macrophages could mediate damage to the phagosomal membrane in a manner similar to PLB1(De Leon Rodriguez et al., 2018), and may contribute for the association the alpha mating locus with virulence in *C. neoformans*. These hypotheses remain to be tested. Another protein potentially associated with virulence present in macrophage-like cells was Tra nsaldolase (Tal1). While the function of Tal1 in *C. neoformans* virulence has not been widely studied, there is evidence for its role in resistance to nitric oxide, which could contribute to fungal cell survival in macrophage-like cells (Missall et al., 2006). We note that a large proportion of the peptides identified belong to so called ‘hypothetical proteins’ identified in the *C. neoformans* genome. Our data strongly suggests that these hypothetical proteins are expressed during infection and *in vivo*, providing fertile ground for new investigations into their function in the physiology and virulence of *C. neoformans*.

Among the proteins found in macrophage-like cells infected with live *C. neoformans* was the ATP-binding cassette transporter (ABC transporters) Yor1. Yor1 and other ABC transporters have a variety of roles in other organisms, but one overarching theme is the translocation of solutes across membranes which can include polysaccharides, peptides, ions and lipids (Cangelosi et al., 1990; Davidson and Chen, 2004; Decottignies et al., 1998; Henderson and Payne, 1994; Kihara and Igarashi, 2004; Kumari et al., 2021; Nishida and Tsubaki, 2017). ABC transporters are characterized by multiple transmembrane domains and an ATPase-domain; in microbes, this family has an enormous array of functions ranging from iron or nutrient import (Henderson and Payne, 1994; Kumari et al., 2021) to the export of virulence factors such as capsular polysaccharide (Orsi et al., 2009). In *S. cerevisiae*, Yor1 influences the cell membrane composition by translocating phosphatidylethanolamine outwards (Khakhina et al., 2015), and is necessary in efflux of beauvericin, a microbial product which potentiates activity of antifungal drugs (Shekhar-Guturja et al., 2016). In *Candida albicans*, Yor1 is required for efflux of geldanamycin, a Hsp90 inhibitor (Hossain et al., 2021), and has been implicated in antifungal resistance in *C. lusitaniae* (Reboutier et al., 2009), but its role in fungal homeostasis has not been studied. Since the role of Yor1 in *C. neoformans* physiology was unknown, we performed some exploratory assays with *yor1*Δ strains, including interaction with mammalian cells. We observed no difference in growth rates in microbiological media, but detected alterations in. capsular size in minimal media and a decrease in melanin secretion in *yor1*Δ deletion strains, hinting at the possibility that it may be involved in export of virulence factors. Analysis of the frequency of non-lytic exocytosis for *C. neoformans* strains deficient in Yor1 revealed reduced rates of fungal cells exiting macrophage-like cells. Yor1 is involved in transport of still unknown moieties that affect proteins or lipids needed for secretion of virulence factors, which overall decrease melanin secretion and capacity to provoke non-lytic exocytosis. This is linked with reduced capacity to kill wax moth larvae.

In summary, we report a new method to investigate proteins secreted by microbes ingested by macrophages. We reason that this protocol may be extended to other systems and even *in vivo* in animal models. We show that ingestion of live and dead *C. neoformans* cells by macrophage-like cells results in release of different protein sets, consistent with active release and digestion of fungal cells, respectively. The finding that some of these proteins modify the course of *C. neoformans* intracellular pathogenesis is consistent with recent findings that fungal cells actively modulate some processes such as non-lytic exocytosis and cell-to-cell transfer (dragotcytosis) (Dragotakes et al., 2019). The discovery that many of the proteins putatively secreted into macrophage-like cells are poorly characterized provides a rich trove of new research avenues that could reveal fundamentally new processes in intracellular *C. neoformans* pathogenesis.

## Materials and Methods

### Strains

H99 (Serotype A) fungal cells from frozen stocks (10% glycerol) were plated on rich Sabouraud agar plates with single colonies subsequently picked and maintained in Difco™ Sabouraud Dextrose Broth (BD, Sparks, MD) at 30° C with agitation for 18 hr. Our initial screens were conducted with a mutant library generated by Drs. Hiten Madhani and Suzanne Noble at University of California San Francisco and made publicly available at Fungal Genetic Stock Center. The library included parental H99 and approximately 2,000 gene knockouts (Liu et al., 2008). The strain used for *de novo* mutant generation is a derivative of H99, H99-FOA with a URA5 selectable marker (Edman and Kwon-Chung, 1990).

### *C. neoformans* protein isolation

Cryptococcal protein isolation was accomplished by adding equal parts prewashed (with ice cold PBS with protease inhibitors), H99 and 0.5 mm Zirconia/Silica Beads (Biospec, Bartlesville, OK) to a microcentrifuge tube and vortexing at max speed for 4 cycles of 5 minutes agitation, 2 minutes on ice. The lysate was centrifuged, 4000 × g at 4° C for 5 minutes and the supernatant was removed and sequentially centrifuged 3 times to remove cellular debris, intact *C. neoformans* and beads while leaving proteins in the resulting supernatant.

### Macrophage-like cell lines

J774.16 is a murine macrophage-like cell line. From frozen stock, J774.16 cells were washed several times to remove freezing media and plated on non-cell culture treated dishes. They are maintained in DMEM (Gibco), 10% NCTC-109 medium (Gibco), 10% heat-inactivated FBS (Atlanta Biologicals), and 1% nonessential amino acids, at 37° C with 9.5% CO_2_. For infection, macrophage-like cells from anywhere between passages 4 and 10 are plated at ∼60% confluency on 150 mm non-cell culture treated dishes (Corning^®^ #430597) and activated with 150 U/mL interferon gamma and 10 ng/mL lipopolysaccharide overnight. Infection is accomplished under the same conditions with the addition of 10 µg/mL of 18B7 (an opsonic monoclonal antibody against *C. neoformans’* polysaccharide glucuronoxylomannan capsule) for 2 h (Casadevall et al., 1998).

Bone marrow derived macrophage isolation, differentiation and infection. All animal experiments were approved by Johns Hopkins University IACUC under protocol number MO18H152. Primary bone marrow derived macrophage cells were used to study non-lytic exocytosis. These cells were isolated from 6-8-week-old C57Bl/6 female mice. Bone marrow derived macrophages (BMDM) were differentiated from bone marrow at 37° C with 9.5% CO_2_ for 7 days. BMDM media consists of DMEM, 20% L929 conditioned media, 10% FBS, 1% non-essential amino acids, 1% GIBCO™ GlutaMAX™, 1% HEPES, 1% penicillin-streptomycin and 0.1% β-mercaptoethanol. After 7 days, the differentiated BMDMs were detached using CellStripper™ (Corning, Corning, NY) and 10^4^ BMDMs were plated on MatTek™ dishes which have a central glass coverslip insert in the center of a hollowed polystyrene plate and allows for time-lapse microscopy. Fungal strains were counted and opsonized with 18B7 at an MOI of 3:1 for 2 h at 37° C and 9.5% CO_2_. After 2 h infection, the plates were washed 3 times with BMDM media for subsequent 24 h time-lapse microscopy at 37° C and 9.5% CO_2_ and non-lytic exocytosis analysis.

### Non-lytic exocytosis analysis

Wild type or *yor1*Δ strains were resuspended and opsonized with 18B7 (monoclonal antibody against the capsular polysaccharide of *C. neoformans*) to infect at MOI of 3:1 for 2 h at 37° C and 9.5% CO_2_. After 2 h infection, the plates were washed 3 times with BMDM media for subsequent 24 h time-lapse microscopy at 37° C and 9.5% CO_2_ and non-lytic exocytosis analysis. Infected macrophage-like cells were tracked over the course of 24 h and non-lytic / lytic exocytosis was assessed against infected macrophage-like cells that remained mobile and infected through 24 h.

### Protein isolation from *C. neoformans*-infected macrophage-like cells

10 cm plates with monolayers of approximately 2.0 × 10^7^ *C. neoformans* infected J774A.16 macrophage-like cells or bone marrow derived macrophages (BMDM), were rinsed with warm PBS to remove non-internalized fungus and unattached dead macrophages. After scraping the remaining infected macrophage-like cells from the plate, they were centrifuged at 1000 × g for 5 minutes at 4° C to pellet the infected macrophage-like cells. The supernatant was discarded and the pellets were resuspended in ice-cold water containing cOmplete™ protease inhibitors (Roche). Each sample was subsequently passed 10 times through a 26-gauge needle to shear the host macrophage-like cells while leaving the fungi intact. Visual inspection under a microscope confirmed macrophage lysis while fungal counts confirmed the viability of *C. neoformans* through the lysis procedure. Most *C. neoformans* and cellular debris was removed from the lysate by an initial centrifugation of 3,000 × g for 10 minutes at 4° C. Two subsequent 8,000 × g centrifugations of the resulting supernatants for 10 minutes at 4° C removed all cellular debris and remaining *C. neoformans*. Proteins were concentrated in a Savant Speed Vac Concentrator. Total proteins were quantified using Pierce™ BCA Protein Assay Kit (ThermoFisher Scientific). 50 µg of each total lysate (mixture of murine and cryptococcal proteins) from the protein isolation step was run on a 12% NuPAGE™ tris-acetate gel (Thermo Fisher Scientific) two centimeters into the gel prior to Coomassie R250 staining and water de-staining for subsequent proteomic analysis.

### Proteomics

Resulting gel lanes containing a mixture of fungal and mammalian proteins were excised and fractioned into five sections prior to in-gel protein digestion and were analyzed on an Orbitrap Fusion™ Tribrid™ (Thermo Scientific) mass spectrometer at the Herbert Irving Comprehensive Cancer Center (Columbia University Medical Center) proteomics facility by Dr. Emily Chen. Peptide data was aligned and filtered using murine and cryptococcal databases. Mass spectrometry proteomics data have been deposited to the ProteomeXchange Consortium via the PRIDE (PubMed ID: 30395289) partner repository with the dataset identifier PXD024951.

### Generation of *de novo* deletion strains

Flanking regions of 1 kb to the genomic sequence of *YOR1* (CNAG_03503) and the selection marker, *URA5*, were amplified by PCR to contain overlapping nucleotides for a 2-step fusion PCR. For step 1, *YOR1* was amplified from *C. neoformans* genomic DNA as a template; *URA5* was amplified from plasmid DNA containing the *C. neoformans* actin promoter and the coding sequence for *URA5* (Sup. Table 2). All primers contained annealing overhangs for subsequent fusion PCR. After gel purification of all PCR products, the second step of fusion PCR yielded a complete sequence including the 5’ 1000 bp region of genomic *YOR1, URA5* selectable marker and the 3’ 1000 bp region of genomic *YOR1*. This combined fusion PCR product was purified in preparation for biolistic particle delivery.

### Biolistic Particle Delivery and screening

The Bio-Rad PDS-1000 Biolistic Particle Delivery system was used to replace *YOR1* in H99-FOA *C. neoformans*. H99-FOA is an auxotrophic mutant lacking the ability to grow without supplemental uracil and this strain was selected for its ease in knockout generation. Inserting *URA5* as a selection marker allows for transformants to grow without supplemental uracil. H99-FOA was grown to stationary phase overnight in rich YPD media at 30° C. After plating a lawn on 1 M sorbitol-YPD plates, the plates were dried in a sterile hood. The DNA preparation involves coating 10 μl of .6 μm Gold microcarriers (1652262 Bio-Rad, Hercules, CA) with 1μg of purified DNA by gently vortexing DNA, 10 μl 2.5 M CaCl_2_ and 2 μl 1 M spermidine-free base before washing with 100% EtOH. 10 μl of the DNA-coated microcarrier beads dried on 2.5 cm microcarrier discs (1652335 Bio-Rad, Hercules, CA). The manufacturer’s protocol for operation of the PDS-1000 system was followed. Transformed plates were allowed to recover overnight at room temperature before subsequent streaking on rich media plates where isolates were selected. For screening of successful transformants, cells from 50 colonies were lifted and pooled in groups of 10. Successful replacement of *YOR1* was tested using colony PCR to amplify the *YOR1* locus. Successful transformants contained a shorter product with *URA5* (1.3 kb) replacing the endogenous *YOR1* gene (5 kb). Three de novo Δ*yor1* mutants were isolated were confirmed with single colony PCR and named *yor1*Δa, *yor1*Δb and *yor1*Δc.

### Characterization of *yor1*Δ Mutants

To assess the cell body and capsule size of *yor1*Δ mutants versus wildtype H99, both strains were grown to stationary phase at 72 h at 30° C in minimal media (MM) composed of 15 mM dextrose, 10 mM MgSO_4_, 29.4 mM KH_2_PO_4_, 13 mM glycine, and 3 μM thiamine-HCL with constant agitation. Liquid cultures were centrifuged at 4,000 × *g* for 15 min to pellet the cells. The supernatant was saved for vesicle purification, and the pellet was resuspended in PBS for subsequent India ink negative stain imaging. Images were captured on an Olympus IX 70 microscope (Olympus America Inc. Melville, NY) to further compare mutants with wildtype H99. Cell body and capsule radius were measured using QCA, the custom cell body and capsule measurement application (Dragotakes et al., 2019). The Bioscreen C (Growth Curves USA) was used to evaluate any growth defects in *yor1*Δ strain versus wildtype H99. Cultures were grown Sabouraud rich media overnight from frozen stock. 24 h later, they were counted via hemocytometer before inoculating either Sabouraud or minimal media at a concentration of 5 × 10^5^ before 10-fold serial diluting to 5 × 10^2^ per ml. 200 μl of each culture was plated in triplicate on HC-2 honeycomb plates (Growth Curves USA) yielding a starting inoculum of 10^5^ down to 10^2^ cells per 200 μl well. The plates were run with constant agitation at 37° C for 72 h with a wideband (OD_420-580_) measurement taken every 15 mins. Alternatively, growth curves were performed by seeding 10^3^ cells/mL in 2 mL media (SAB or MM) in 12-well tissue culture dishes. Cultures were then incubated for 96 h at 30 °C with OD600 measurements taken every 2 h on a BioTek Epoch microplate spectrophotometer.

### Melanization assay

*C. neoformans* strains were grown overnight in YPD broth at 30□C until they were in stationary phase. Cultures were washed twice in PBS, and inoculated into Minimal Media at 10^6^ cells/ml with 1 mM L-3,4-dihydroxyphenylalanine (L-DOPA). Cells were grown at 30? and monitored daily for pigment formation. Cultures were imaged after 3 days.

### Infection of *Galleria mellonella*

Final instar *Galleria mellonella* larvae were obtained from Vanderhorst Wholesale Inc, St. Marys, Ohio, USA. Healthy cream-colored larvae roughly between 175 and 225 mg were sorted and separated into groups of equal numbers. C. neoformans strains were grown overnight in 1 ml of YPD broth until the culture was in stationary phase. Larvae were then infected with 10 µl of 10^7^ C. neoformans cells/ml in a PBS suspension. Survival was measured daily over the course of 10 days, with survival assayed by observing larval and pupal movement following stimulation with a pipette tip. The Cox Mixed Effects and Hazard Ratio calculations were performed using R for R 4.0.2 GUI 1.72 for Mac OS at https://www.r-project.org/ (R Core Team, 2020) and the *coxme* package, version 2.2-16 (Therneau, 2020).

### Measurements of phagolysosomal pH

Phagolysosomal pH was measured by an established ratiometric fluorescence technique, as described in previous studies(Dragotakes et al., 2020; Fu et al., 2018). Briefly, BMDM were seeded on glass coverslips in 24-well tissue culture plates at a density of 1.25 × 10^5^ cells/well and activated overnight with IFNγ (100 U/mL) and LPS (500 ng/mL). *C. neoformans* particles were opsonized with 18B7 monoclonal antibody, which was previously conjugated to Oregon Green 488, at a final concentration of 10 µg / mL and added to the activated macrophages at MOI 1:1. The plates were centrifuged at 350 x *g* for 1 min to synchronize fungal cell adherence and ingestion by macrophages. After 2 h, media in each well was replaced twice with an equivalent volume of HBSS and the coverslip placed upside down on a MatTek Petri dish with HBSS in the microwell for imaging on an upright scope (Olympus AX70). Ratiometric fluorescence measurements focusing on ingested intracellular *C. neoformans* cells were made using 440 nm and 488 nm excitation and 520 nm emission. Images were analyzed using Metafluor Fluorescence Ratio Imaging Software to calculate fluorescence ratios. The pH of the phagosome was calculated from interpolation of a standard curve; the standard curve was generated by incubating infected macrophages with HBSS buffer with 10 µM nigericin and of known pH ranging from pH 3 to 7.

### Non-lytic exocytosis assays

The Axiovert 200 (Zeiss, Oberkochen, Germany) and a Hamamatsu ORCA ER cool charged-coupled device (CCD) with a heated chamber and supplemental CO_2_ were used for all time-lapse microscopy. Time-lapse microscopy was done at 10x bright-field with an image captured every 4 min for 24 h to assess host-pathogen interaction. Time-lapse was compiled in ImageJ and 50 to 100 infected macrophage-like cells were tracked through the course of 24 h. Type I non-lytic exocytosis was counted when all yeasts were expelled from the host macrophage with both remaining intact. Type II was counted when part of the fungal burden was expelled while a portion remained within the host macrophage. Type III, or cell-to-cell transfer, was counted when an infected BMDM would pass one or more yeast directly to another BMDM. Infected macrophage-like cells that underwent lytic exocytosis were marked as lysed.

## Supporting information

Sup Table 1. C. neoformans proteins identified in J774.16 macrophage infected with H99.

## Bioinformatics analysis

We performed bioinformatics analysis using SignalP (http://www.cbs.dtu.dk/services/SignalP/)(Almagro Armenteros et al., 2019), PrediSi (http://www.predisi.de/), and Phobius (https://phobius.sbc.su.se/)(Kall et al., 2007). Structure prediction was performed using Phyre2 (http://www.sbg.bio.ic.ac.uk/phyre2/html/page.cgi?id=index) (Kelley et al., 2015). Results are displayed in Supplemental Table 1.

## Funding and Acknowledgments

EHJ, QD, LSR, DQS, AD, AJ, CC and AC were supported by NIH grants to AC [AI052733-16, AI152078-01, and HL059842-19]. CC is now supported by grants to the MRC Centre for Medical Mycology (Exeter, UK). This work was supported, in part, by the Intramural Research Program of the National Institutes of Health [AI001123 and AI001124 to PRW].

We acknowledge the help of Dr. Emily Chen, from Herbert Irving Comprehensive Cancer Center (Columbia University Medical Center) proteomics facility.

## Supplemental Information

**Supplemental Table 1**. *C. neoformans* proteins during infection of murine macrophages.

---

**Supplementary Figure 1.**
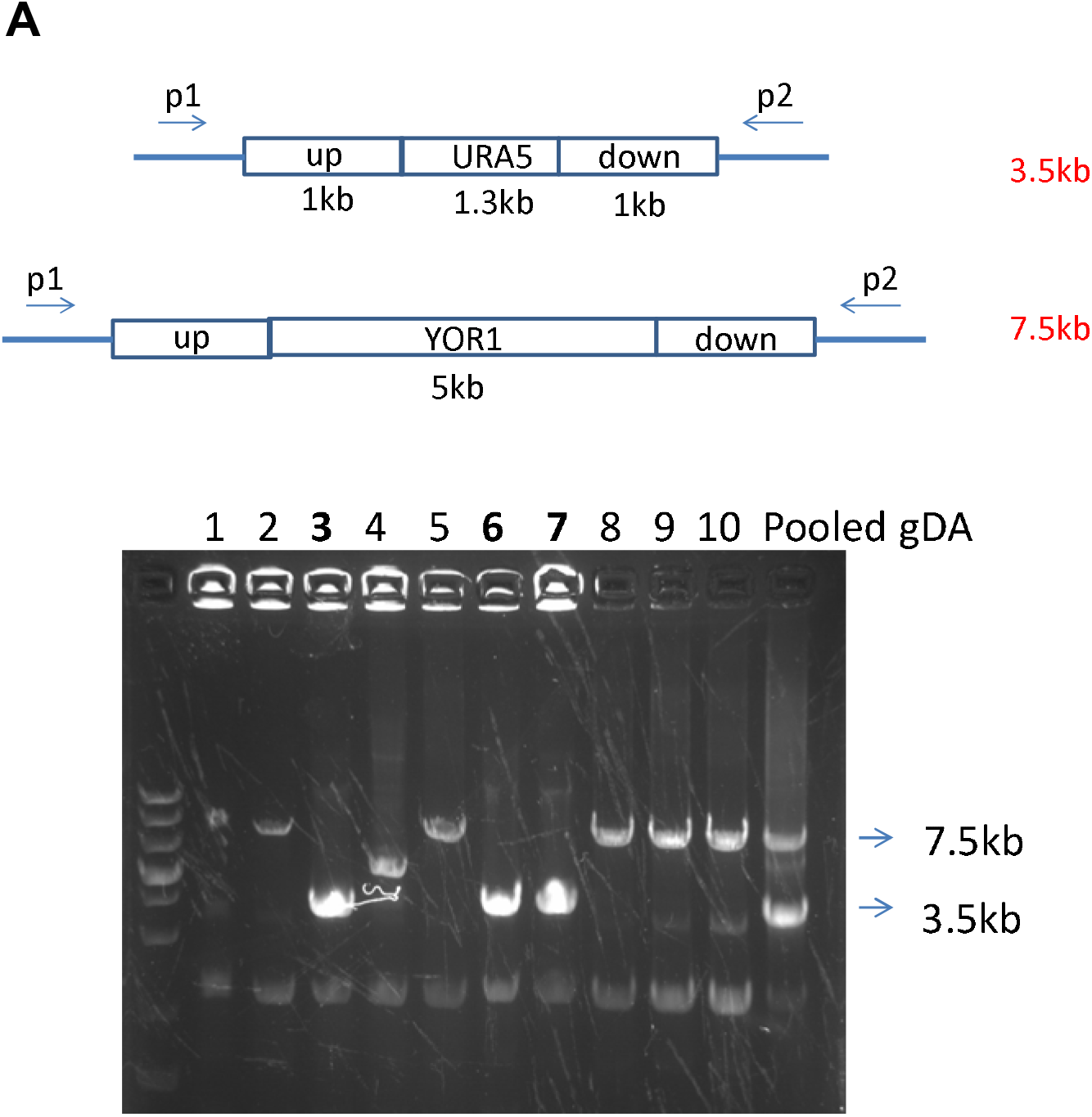
Screening of *de novo yor1*Δ strains. Successful deletion was detected with colony PCR. (A) Diagram of primer locations for colony PCR. (B) Insert size of individual clones, showing lanes 3,6,7 with *Yor1* deleted.

**Supplementary Figure 2.**
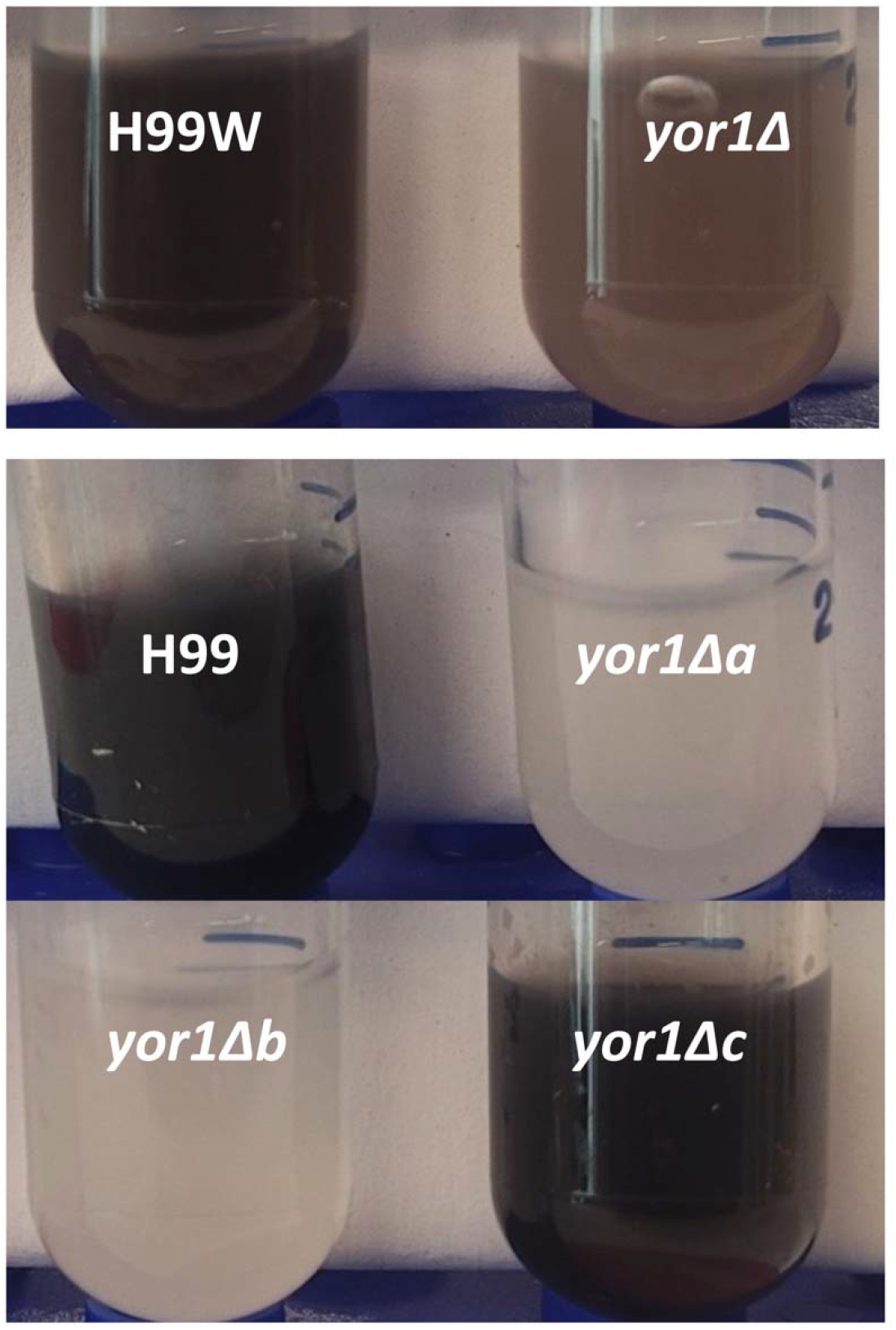
Melanin secretion is affected by *Yor1* deletion. Strains were grown in minimal media with L-DOPA for several days. Images are representative image of cultures from 2-3 independent biological replicates.

**Supplementary Figure 3.**
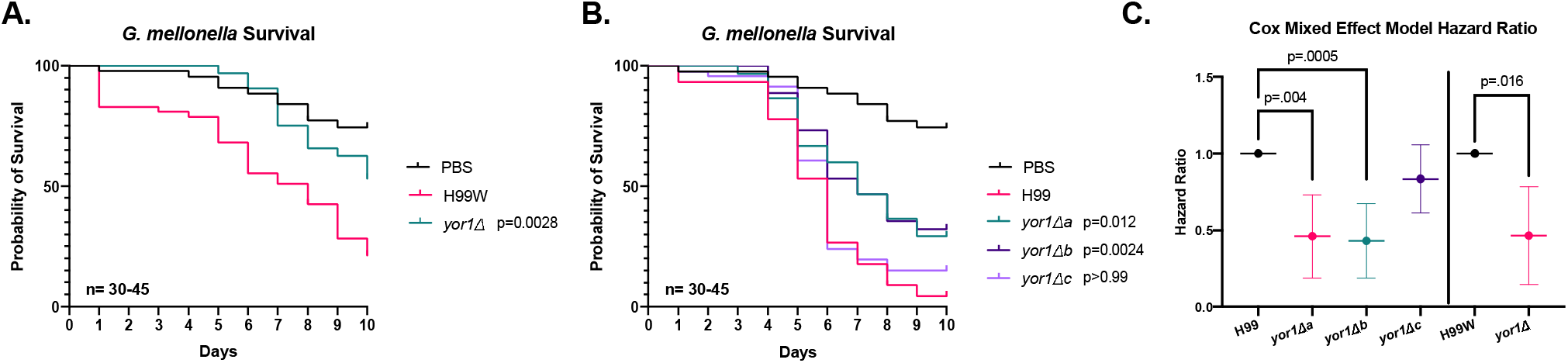
*YOR1* is required for Virulence in *Galleria mellonella*. Deletion of *YOR1* reduces virulence of *C. neoformans* during infection of *G. mellonella*. (**A**) Infection of 10^5^ CFU/larvae with the *yor1Δ* compared to its parental strain (H99W). (**B)** Infection of 10^5^ CFU/larvae with the *yor1Δ*a-c compared to its parental strain (H99). (**C)** The hazard ratios calculated using a Cox Mixed Effects model to account for random variables between replicates show similar reductions in Hazard Ratios for the 2008 library *yor1Δ* strain, and for the independently generated *yor1Δ*a and *yor1Δ* mutants relative to their respective parental strains Infections in panels **A, B** were performed simultaneously, with the same uninfected control group (PBS), and were separated to facilitate visualization. Survival statistics in **A, B** were performed using the Mantel-Cox Log Rank test, with a Bonferroni correction for four comparisons. Survival experiments represent 2 to 3 independent experiments, with each group containing 15 larvae in each independent experiment.

**Supplementary Figure 4.**
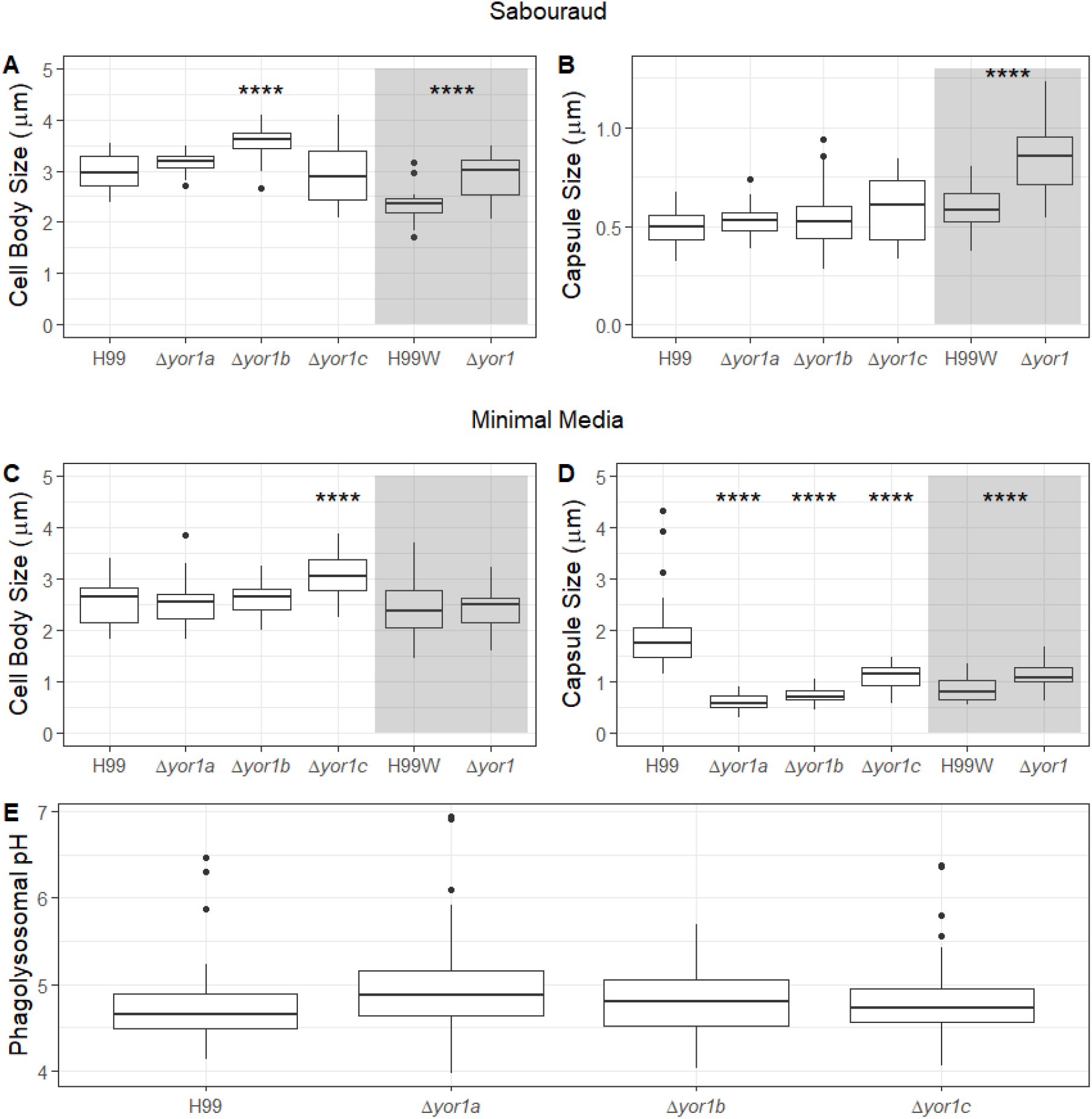
*Yor1* deletion affects capsule production of *C. neoformans* grown in minimal media, albeit with contribution of strain background. Cell body and capsule radius measurements of *yor1*Δ strains compared to the parental H99 strain. (A) Cell body radius measurements of strains after 2 d culturing in Sabouraud media. (B) Capsule radius measurements after 2 d culture in Sabouraud media. (C) Cell body radius measurements of strains after 2 d culturing in minmal media. (D) Capsule radius measurements of strains after culturing 2 d in minimal media. (E) Measurements of pH of *C. neoformans* containing phagolysosomes 2 h post ingestion by **J774**.**16** **** signifies *P* < 0.0001 via ANOVA with Dunnett’s multiple comparisons to parental H99 strain.

**Supplementary Figure 5.**
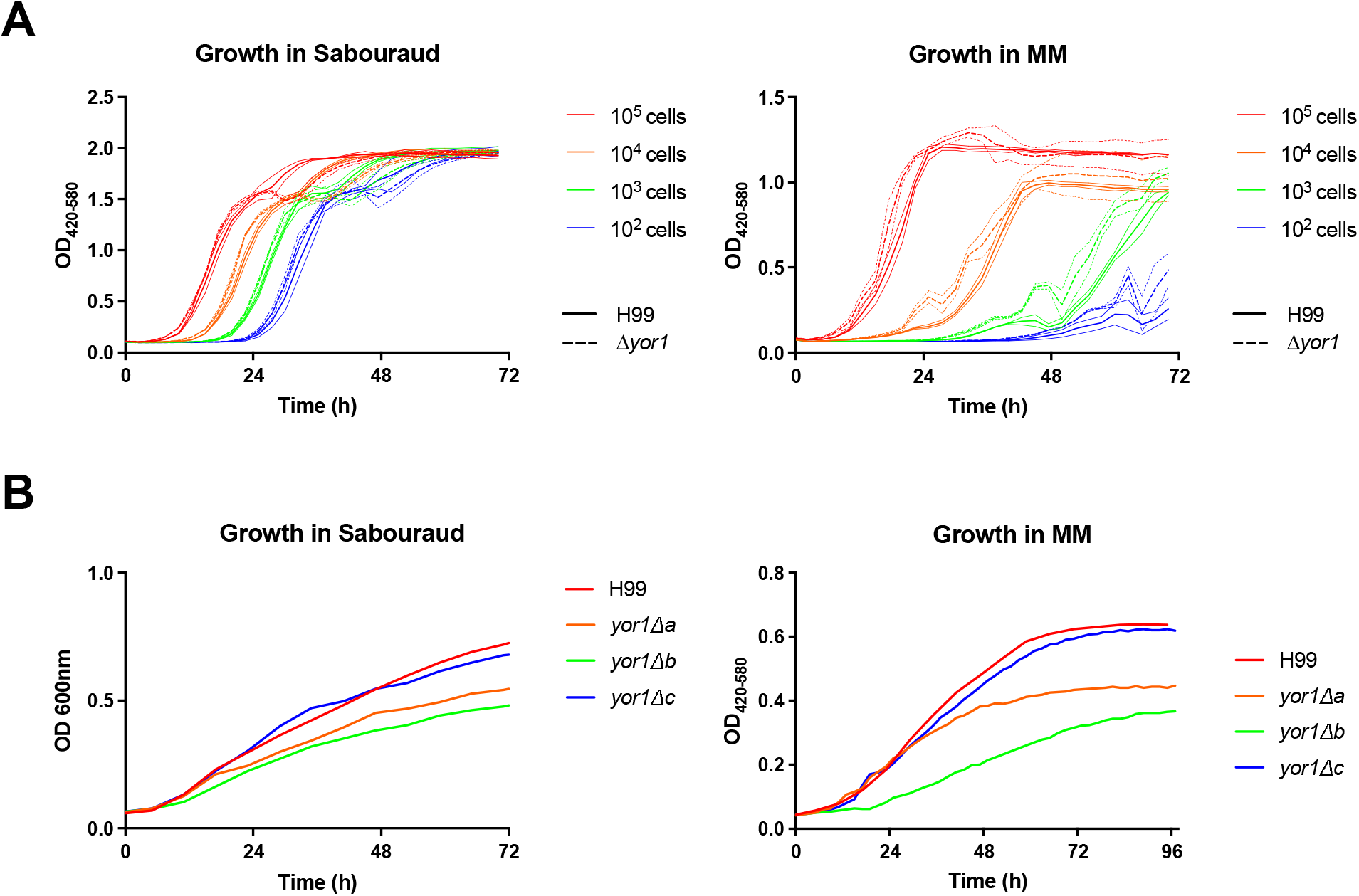
No detectable growth defect in absence of *yor1* Δ. **(A)**To measure growth defects in wildtype versus *yor1* Δ strains, a 10-fold dilution series of starting inoculum was run in the Bioscreen C at 37° C with wideband (OD_420-580_) measurements taken every 15 mins for 72 h in both rich Sabouraud broth and minimal media (MM) broth. Represented is every 9^th^ data point, and standard deviation shown as outside thinner lines. Experiment performed once in experimental triplicates. (B) Growth curves were performed in a BioTek Epoch microplate spectrophotometer, with measurements taken every 2h for 96h at 30°C. Represented is every 2^nd^ data point, Experiment was performed once.

